# No single, stable 3D representation can explain pointing biases in a spatial updating task

**DOI:** 10.1101/390088

**Authors:** Jenny Vuong, Andrew W. Fitzgibbon, Andrew Glennerster

## Abstract

People are able to keep track of objects as they navigate through space, even when objects are out of sight. This requires some kind of representation of the scene and of the observer’s location but the form this representation might take is debated. We tested the accuracy and reliability of observers’ estimates of the visual direction of previously-viewed targets. Participants viewed 4 objects from one location, with binocular vision and small head movements giving information about the 3D locations of the objects. Without any further sight of the targets, participants walked to another location and pointed towards them. All the conditions were tested in an immersive virtual environment and some were also carried out in a real scene. Participants made large, consistent pointing errors that are poorly explained by any consistent 3D representation. Instead, if a 3D representation is to account for the data it would need to be one where the target boxes were squashed, almost into a plane, quite far away from the true location of the boxes and in different places depending on the orientation of the obscuring wall at the moment the participant points. In short, our data show that the mechanisms for updating visual direction of unseen targets are not based on a stable 3D model of the scene, even a distorted one.

## 1. Introduction

If a moving observer is to keep track of the location of objects they have seen earlier but which are currently out of view, they must store some kind of representation of the scene and update their location and orientation within that representation. There is no consensus on how this might be done in humans. One possibility is that the representation avoids using 3D coordinates and instead relies on a series of stored sensory states connected by actions (‘view-based’), as has been proposed in simple animals such as bees or wasps [1, 2, 3], humans [4, 5, 6] and deep neural networks [7, 8]. Alternatively, the representation might be based on 3D coordinate frames that are stable in world-, body- or head-centered frames [9, 10], possibly based on ‘grid’ cells in entorhinal cortex [11, 12] or ‘place’ cells in the hippocampus [13, 14].

One key difference between these approaches is the extent to which the observer’s task is incorporated in the representation. For 3D coordinate-based representations of a scene, task is irrelevant and, by definition, the underlying representation remains constant however it is interrogated. Other representations do not have this constraint. Interestingly, in new approaches to visual scene representation using reinforcement learning and deep neural networks, the task and the environment are inextricably linked in the learned representation [15, 7]. This task-dependency is one of the key determinants we explore in our experiment on spatial updating.

If the brain has access to a 3D model of the scene and the observer’s location in the same coordinate frame then, in theory, spatial updating is a straight-forward matter of geometry. It is harder to see how it could be done in a view-based framework. People can imagine what will happen when they move [16, 17, 18] although they often do so with very large errors [19, 20, 21, 22, 23, 24]. In this paper, we examine the accuracy and precision of pointing to targets that were viewed from one location and then not seen again as the observer walked to a new location to point, in order to test the hypothesis that a single 3D reconstruction of the scene, built up when the observer was initially inspecting the scene, can explain observers’ pointing directions. The task is similar to that described in many experiments on spatial updating such as indirect walking to a target [25, 26, 27, 28], a triangle completion task [29, 30], drawing a map of a studied environment including previously viewed objects’ location [31, 19, 32], or viewing a set of objects on a table and then indicating the remembered location of the objects after walking round it or after the table has been rotated [33, 34, 35, 36]. However, none of these studies have compared directly the predictions of a 3D reconstruction model with one that varies according to the location of the observer when they point, as we do here.

Spatial updating has been discussed in relation to both ‘egocentric’ and ‘allocentric’ representations of a scene [37, 38, 39, 40] and, in theory, either or both of these representations could be used in order to point at a target. An ‘egocentric’ model is assumed to encode local orientations and distances of objects relative to the observer [41, 16, 33, 38], while an allocentric model is world-based reflecting the fact that the relative orientation of objects in the representation would not be affected by the observer walking from one location to another [40]. People might use both [37, 17, 39], and Wang and Spelke [38] have emphasised consistency as a useful discriminator between the models. So, for example, disorientating a participant by spinning them on a chair should affect pointing errors to all objects by adding a constant bias if participants use an allocentric representation. The argument that Wang and Spelke [42] make about disorientation conflates two separate issues, one about the origin and axes of a representation (such as ‘allocentric’ or ‘egocentric’) and the other about the internal consistency of a representation. In this paper, we focus on the latter. We ask whether a single consistent, but possibly distorted, 3D reconstruction of the scene could explain the way that people point to previously-viewed targets. In doing so, we concentrate on two potential sources of biases, namely errors in encoding (i) the location of the target boxes as seen from the start zone or maintaining this representation in memory (we lump these together) and (ii) the orientation and position of the observer as they walk. For any model based only on the 3D structure of the scene and the observer’s location within it, these are the two key elements that could contribute to bias in updating a target’s location during self-motion.

## 2. Results

Our stimulus was designed so that participants had access to a rich set of information about the spatial layout of 4 target objects whose position they would have to remember. In either the real world or in a head mounted display, they had a binocular view of the scene and could move their head freely (typically, they moved ± 25 cm). The targets consisted of four different colored boxes that were laid out on one side of the room at about eye height (see Fig. 1a) while, on the other side of the room, there were partitions (referred to from now on as ‘walls’) that obscured the target objects from view once the participant had left the original viewing zone (‘Start zone’), see Figs. 1d and 1e. Most of the experiments were carried out in virtual reality (Fig. 1b), although for one experiment the scene was replicated in a real room (Fig. 1a). After viewing the scene, participants walked to one of a number of pointing zones (the pointing zone was not known in advance) where they pointed multiple times to each of the boxes in a specified order (randomized per trial, see Section S1 for details). Fig. 1 shows the layout of the boxes in a real scene (Fig. 1a), a virtual scene (Fig. 1b) and in plan view (Fig. 1c, shown here for Experiment 1). The obscuring walls are shown on the left with a participant pointing in the real scene. The measure of pointing error used was the signed angle between the target and the ‘shooting direction’ (participants were asked to ‘shoot’ at the target boxes with the pointing device), as illustrated in Fig. 1d. Although not all participants had experience of shooting in video games or similar, the instruction was understood by all and this definition of pointing error gave rise to an unbiased distribution of errors for shots at a visible target (Fig. S3), which was not the case for a considered alternative, namely the direction of the pointing device relative to the cyclopean eye (Fig. 1e). In the first experiment, the participant pointed to each of the target boxes eight times in a pseudo-random order (specified by the experimenter) from one of three pointing zones (shown in a, b and c respectively).

**Figure 1:**
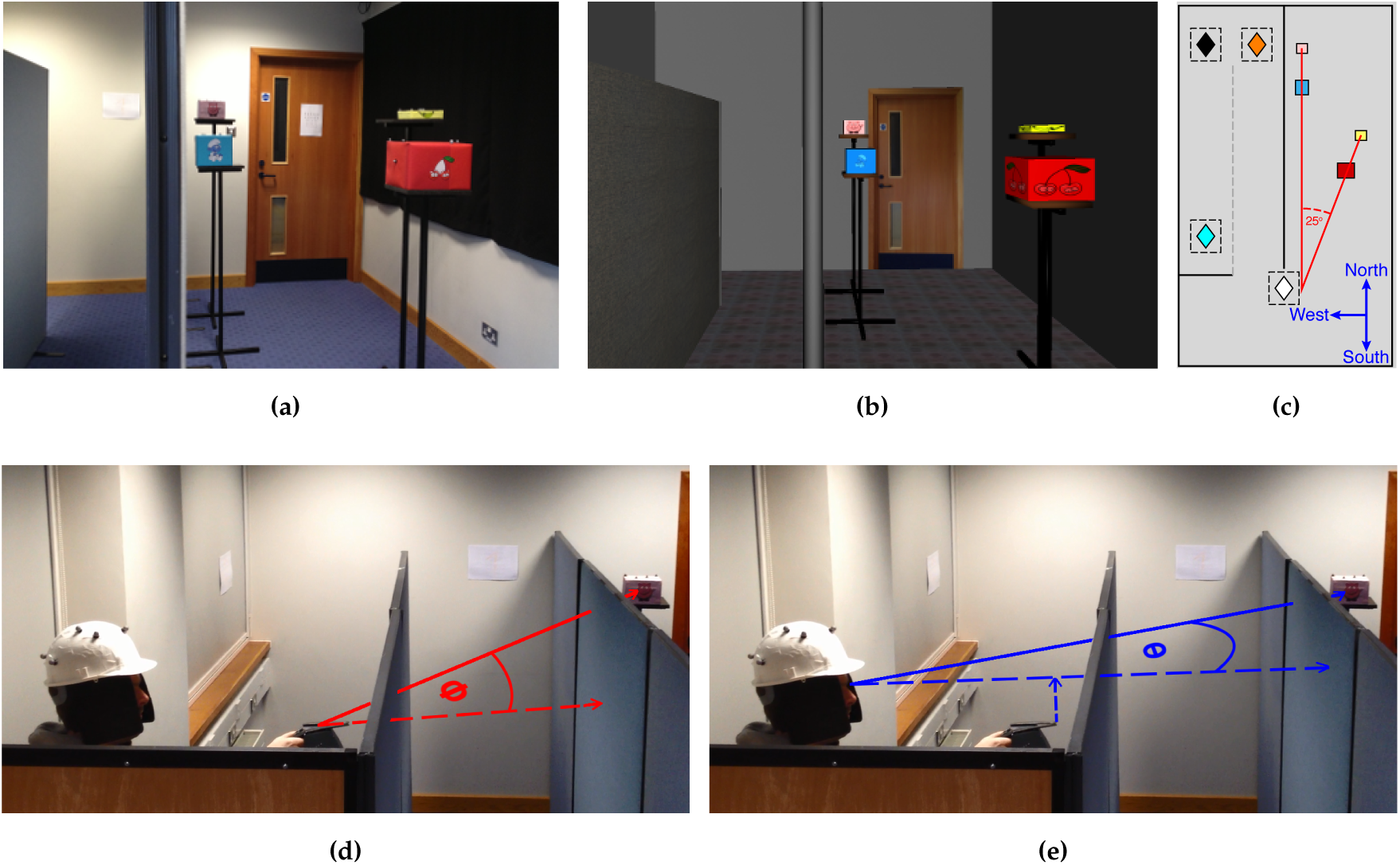
Experimental setup. (a) Real world experiment and (b) in virtual reality. (c) Schematic plan view of one testing layout. Four boxes were arranged such that the blue and the pink, and the red and the yellow boxes lay along two separate visual directions as seen from the start zone (white diamond). From the start zone, the blue and the red box were always closer than the pink and the yellow box. The two visual directions subtended an angle of 25°. This angle was preserved for all box layouts (though distances varied, see Section S1). Pointing to targets was tested at 3 different pointing zones (A = amber, B = black and C = cyan diamond). Black lines indicate positions of walls (in the real world, these were made from partitions). The dashed gray line indicates a wall that disappeared in virtual reality after the participant left the start zone for the ‘direct’ condition. The icon in the corner shows the nominal ‘North’ direction. (d) We defined pointing direction as the direction indicated by the pointing device, labeled here as ϕ. (e) We did not use direction subtended at the cyclopean point (midpoint between the left and the right eye along the interocular axis) and the tip of the pointing device, labeled here as θ.

Figs. 2a–2c illustrate examples of the raw pointing directions in this condition for one participant. It is clear that the participant makes large and consistent errors: For example, they point consistently to the ‘North’ of the targets from the northern pointing zones (A and B) but consistently to the ‘South’ of the targets from the southern zone, C (see Fig. 1c for definition of North). Not only this, some of the geometric features of the scene have been lost. For example, from zone C, the blue and yellow targets are almost co-aligned in reality but the participant points in very different directions to each. As we will see, these features turn out to be highly repeatable, both across participants and in multiple versions of the task.

**Figure 2:**
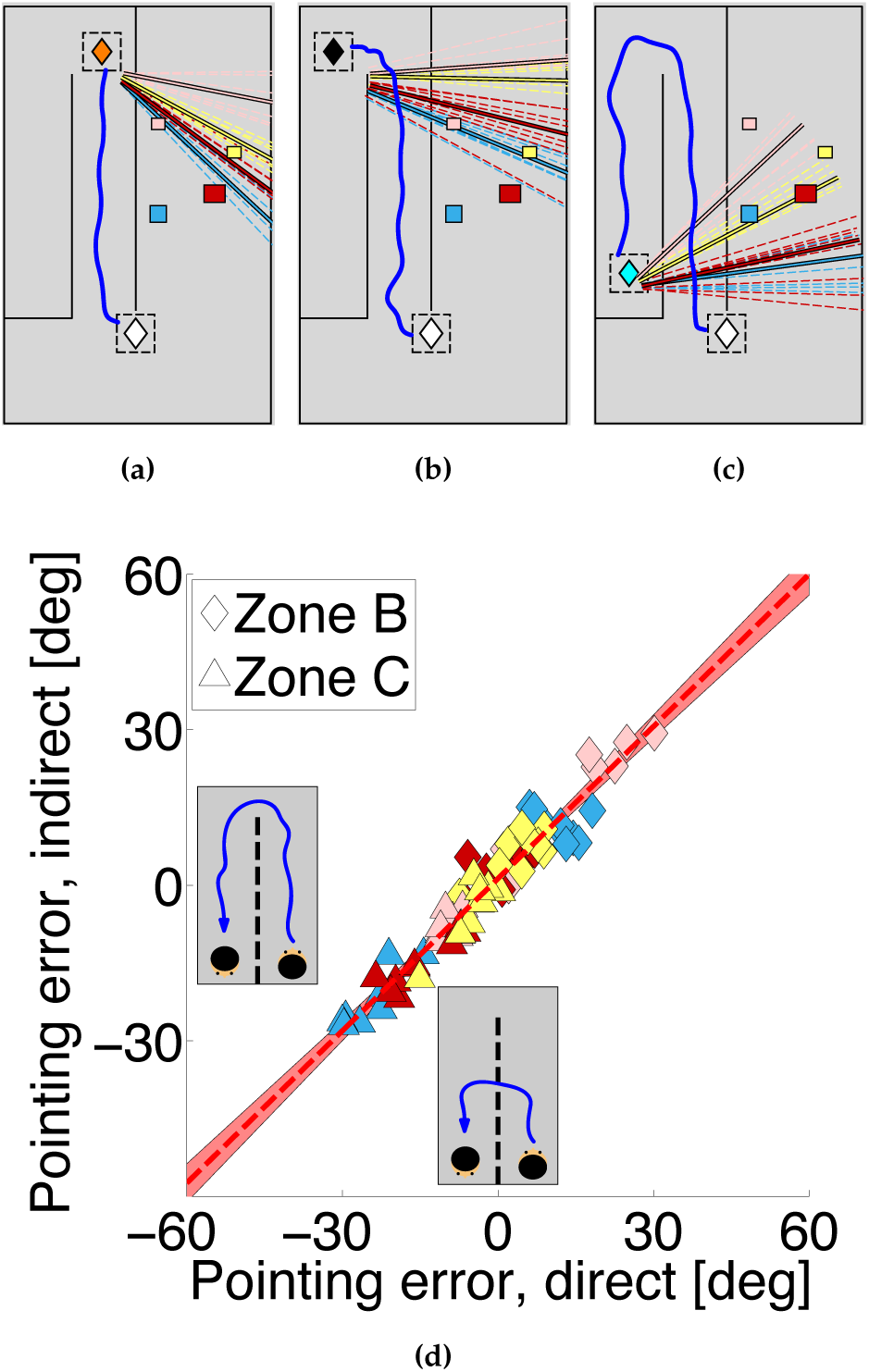
Pointing data from Experiment 1. Plan views in (a–c) show examples of pointing by one participant. The white diamond depicts the center of the start zone. The pointing zone is shown by (a) the amber diamond for zone A, (b) the black diamond for zone B and (c) the cyan diamond for zone C. The blue line shows the walking path from the start zone to the pointing zones. Pointing directions are colour coded according to the target; dashed lines show individual ‘shots’, solid lines indicate the mean for each target. (d) Pointing errors for direct and indirect walking paths to the pointing zones. Each symbol shows mean pointing error for 20 participants and for a given configuration of target boxes. Symbol colours indicate the target box, symbol shape indicates the pointing zone.

For example, Fig. 2d shows the pointing errors gathered in two separate conditions plotted against each other (*Experiment 1*). The data shown are the mean pointing errors for 20 participants shown per target box (symbol colour) for different box layouts and for different pointing zones (symbols shape). Despite the fact that the errors are substantial (from approximately −30° to + 30°), there is a remarkable correlation between the errors in the two experiments (correlation coefficient 0.92, *p* < 0.001, slope 0.98; for individual participant data, see Fig. S2a). The ordinate shows the data from an experiment in virtual reality when the layout of the walls was as shown in Figs. 2a–2c, so that participants had to walk a long indirect route in order to arrive at Zone C, whereas the abscissa shows pointing errors when the experiment was repeated with one of the walls removed (see Fig. 1c) so that participants could walk direct to each of the pointing zones. For zone C especially, this makes a dramatic difference to the path length to get to the pointing zone, so any theory that attributed the pointing errors to accumulation of errors in the estimation of the participant’s location would predict a difference between the data for these two experiments, especially for pointing from Zone C but that is not the case. The data from all the other experiments reported in this paper conform to the same pattern, (see Figs. 3a, 3b, Fig. 4 and Fig. 7), amounting to nine independent replications. We will use the data from this first experiment to build a simple model that predicts the pointing directions in all nine replications and test this model against the alternative hypothesis that the visual system uses a 3D model to generate pointing directions to the unseen targets. For example, Fig. 3a shows that there was also a high correlation between pointing errors when participants repeated exactly the same conditions in a real or a virtual environment (correlation coefficient 0.88, *p* < 0.001), although the range of pointing errors was greater in the virtual room (slope 1.42). Waller and colleagues [43] also found a close match between performance in real and matched virtual environments.

**Figure 3:**
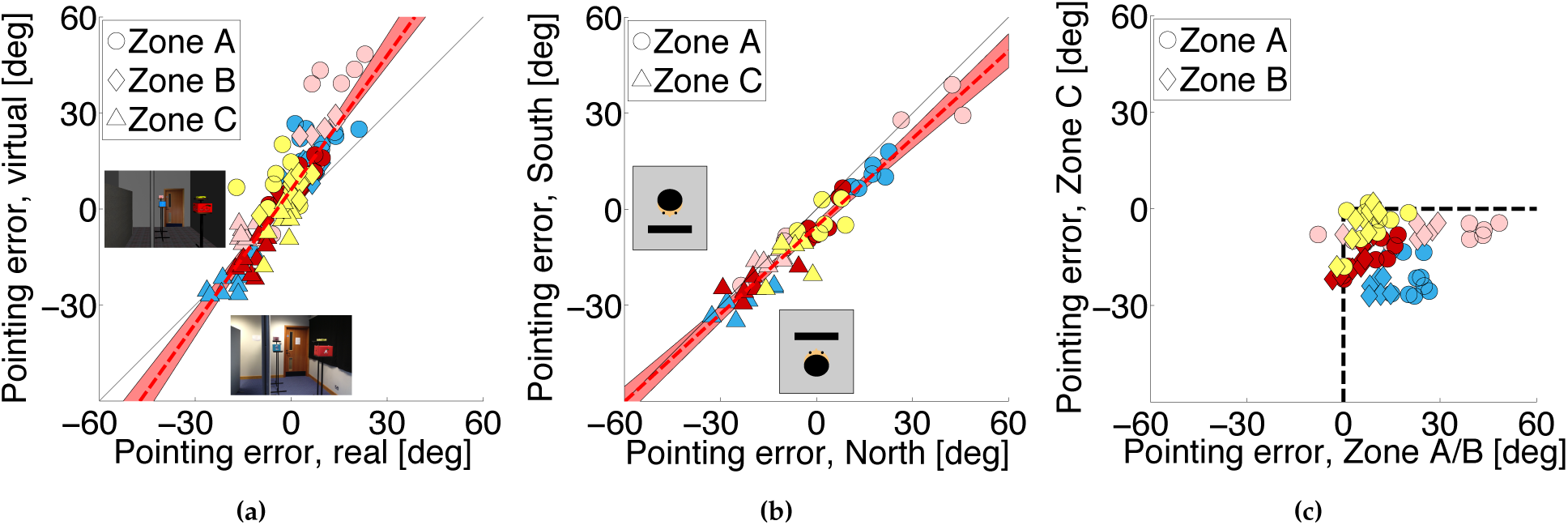
Pointing data from real world and different facing directions. (a) Experiment 1: Pointing errors from real world versus a virtual scene. (b) Experiment 2: Pointing errors for different initial facing directions. (c) Pointing errors for zone C compared to pointing errors for zone A and B (data from Experiment 1 and 2 combined)

**Figure 4:**
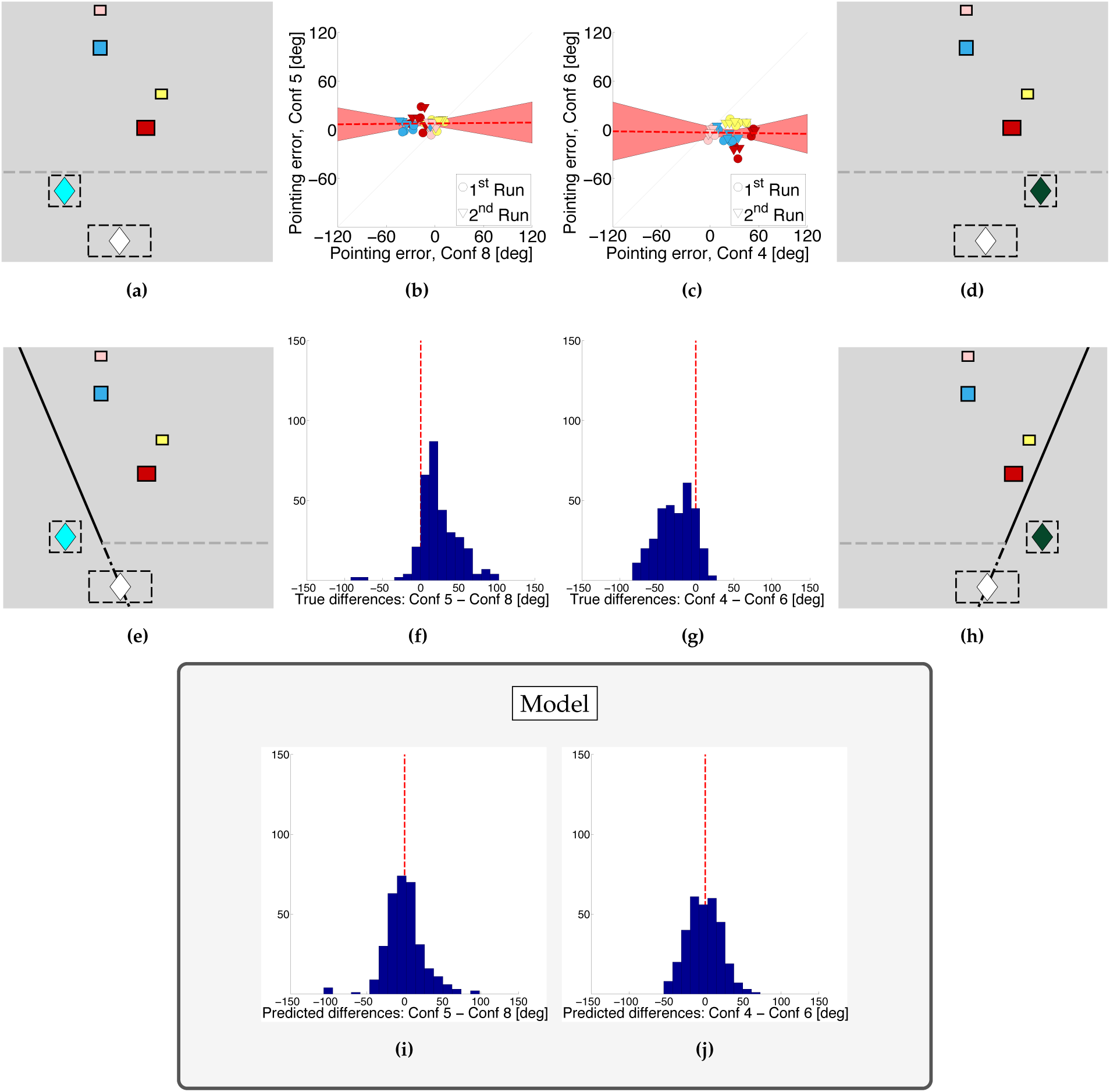
Results of Experiment 4. The same box positions have been tested at the same pointing zones, but differed in the wall orientation (a) versus (e) or (d) versus (h). Dashed gray lines indicate walls that appeared after the participant left the start zone. (b) Pointing errors are plotted for all the paired conditions with an ‘East-West’ wall versus the corresponding condition with a slanted ‘North-West’ wall (pointing from at zone C). (c) Similar comparison but now for a ‘North-East’ wall and pointing from zone E (emerald diamond). (f) Histogram of pointing differences between paired trials for the two different wall orientations pointing from zone C. (g) Same for zone E. (i) and (j) show the same as (f) and (g) respectively, except that they plot pointing differences with respect to the projection-plane model rather than with respect to ground truth.

We found the same pattern of pointing biases in a separate experiment that tested the role of egocentric orientation relative to the targets (*Experiment 2*). Fig. 3b shows a high correlation between pointing errors when participants looked either ‘North’ or ‘South’ in order to view the image that told them which was the next pointing target. In these two versions of the experiment, participants’ rotation to point at the target was quite different, so any model based on egocentric direction at the moment the target was defined would predict a difference, but there was no systematic effect on the pointing errors (correlation coefficient 0.93, *p* < 0.001, slope 0.91). This high repeatability in the face of stimulus changes should be contrasted with the dramatic effect of changing the pointing zone. Fig. 3c shows that, combining data across Experiments 1 and 2, pointing errors when participants were in zones A and B (Fig. 2a–2b) were reliably *anticlockwise* relative to the true target location (positive in Fig. 3c, *M* = 12.1, *SD* = 20.1, *t*(949) = 18.7, *p* < 0.001 in a two-tailed t-test) while, when participants were in zone C (Fig. 2c), the errors were reliably *clockwise* (negative in Fig. 3c, *M* = −12.9, *SD* = 14.7, *t*(479) = −19.2, *p* < 0.001). *Experiment 3*, using pointing zones that were both to the ‘West’ and the ‘East’ of the target boxes showed a similar pattern of biases (see Fig. 7, Fig. S5c and Fig. 8). Another way to summarize the results is that, wherever the pointing zone was, the participants’ pointing directions were somewhere between the true direction of the target and a direction orthogonal to the obscuring wall. Expressed in this way, it is clear that the pointing zone itself may not be the key variable. Rather, it could be the spatial relationship between the target, the screen and the observer at the moment the participant points. In the next experiment, we examined paired conditions in which we kept everything else constant (box layout, start zone and pointing zone) other than the orientation of the obscuring wall.

The results of this experiment (*Experiment 4*) are shown in Fig. 4. Examples of paired stimuli are shown in Fig. 4a/Fig. 4e or Fig. 4d/Fig. 4 h which show the same viewing zone (white diamond), the same pointing zone (cyan or green diamond) and the same box layout but a different obscuring wall (‘East-West’ versus slanted) which appeared after the participant left the viewing zone. It is clear that pointing zone alone is not the only determinant of the biases. Fig. 4b and Fig. 4c plot the pointing error for each condition against the pointing error for its paired stimulus (i.e. changing only the wall orientation) and, unlike the conditions in Fig. 2 and Fig. 3, the data no longer lie close to the line of unity. Instead, as shown in Fig. 4f and Fig. 4 g, the means of the distribution of differences between paired conditions (matched by participant, pointing zone, box layout and target box where only the orientation of the obscuring wall changes) are significantly different from zero (zone C: *M* = 23.5, *SD* = 26.2, *t*(319) = 16.0, *p* < 0.001, zone E: *M* = −27.1, *SD* = 23.8, *t*(319) = −20.3, *p* < 0.001) and shifted in opposite directions for Fig. 4f and Fig. 4 g. This is what one would expect if participants tended to point somewhere between the true target direction and a direction orthogonal to the obscuring wall, just as they did in *Experiment 1* (Fig. 3c). We will return to these paired conditions once we have described a simple model and show that the distribution of pointing directions relative to the *model prediction* (as opposed to the ground truth) is *not* affected by the orientation of the wall (Fig. 4j and Fig. 4i).

## 3. Models

### 3.1. Noisy path integration

The pointing task requires the observer to update an internal representation of their location and orientation. One possible model of our data is that a cumulative error in this updating process is the critical factor explaining the pointing biases (i.e. a noisy-path-integration model). However, as we have already seen from Fig. 2d, participants show a very high correlation between biases obtained when they walk a short distance directly to the target or a longer distance by an indirect route. This would not be expected if pointing biases were caused by cumulative path integration errors, a conclusion supported by modeling. The model assumes that participants maintain an accurate representation of the target location but misestimate their stride length and the rotation of their body on each step. If this noise has zero mean, then the distribution of pointing errors from the final location is also centered on zero (Fig. S4d), but if there is a systematic bias in the estimate of stride length or rotation at each step, then systematic biases in predicted location, and hence in pointing direction, can develop. Fig. 5c and Fig. 5d illustrate this effect for two example trajectories by a participant to zone A (in amber) and zone C (cyan) and, as dotted lines, the model trajectories. Despite the fact that these are the best trajectories fitted to the participants’ pointing data using this model (see Section S4.1 for details), it was always the case that the errors predicted by the model are larger for an indirect path than for a direct path to zone C (Fig. S4b). The participant data does not show this pattern (direct and indirect routes are not significantly different from one another, Fig. 2d). A second problem with this type of model is that it does not make any predictions about the effect of changing the orientation of the obscuring wall (Fig. 4).

**Figure 5:**
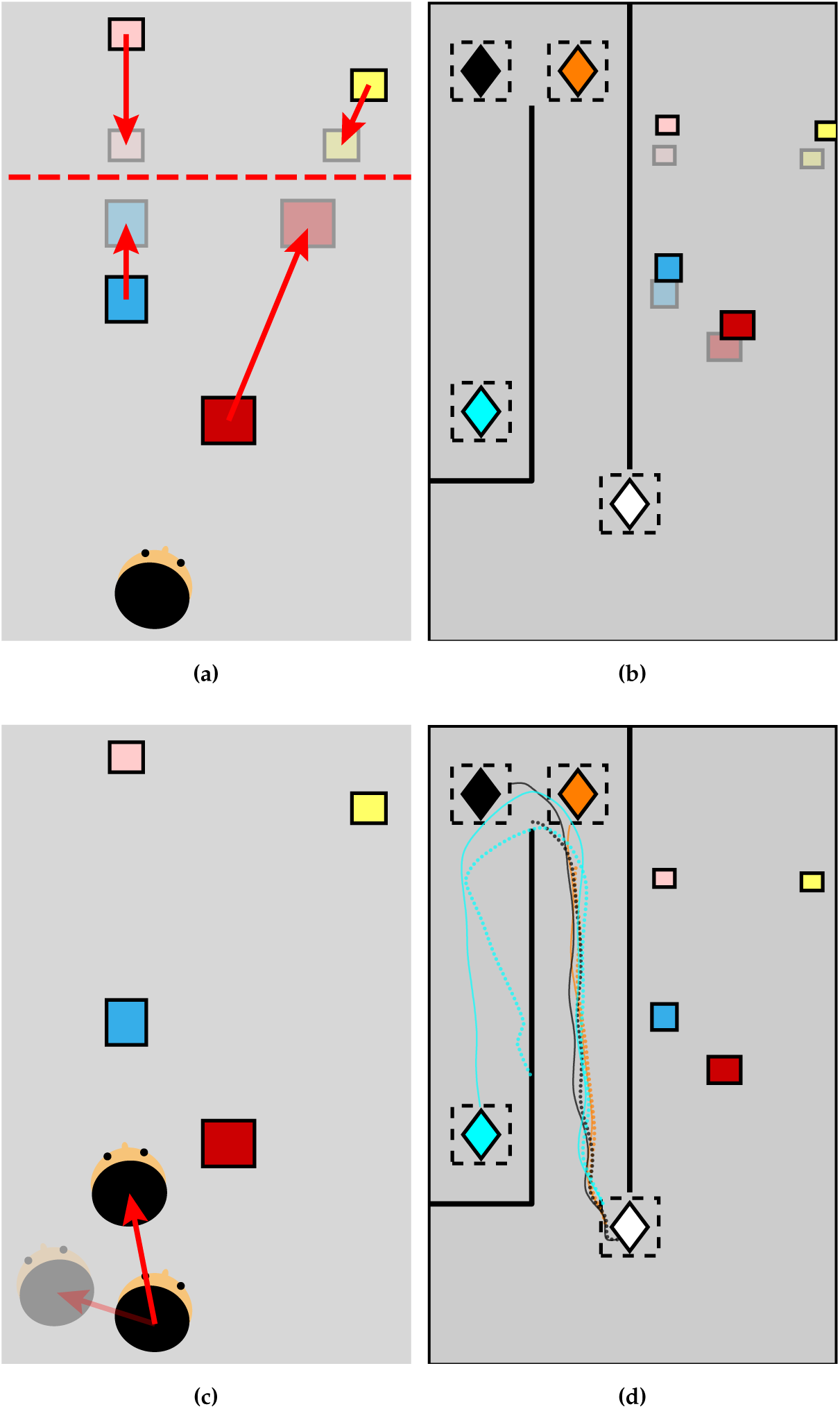
Schematic of abathic-distance-distortion and noisy-path-integration models. (a) The abathic distance is shown by the dashed line and arrows show contraction towards this plane. (b) Example of target box locations shifted under the abathic distance model (opaque rectangles show true box positions). (c) Opaque head shows the observer’s true translation, the faded head shows the assumed translation under the noisy-path-integration model. Free parameters control errors in the estimate of translation and orientation on every step. (d) Solid lines show the walking paths of a participant, dotted lines show the misestimated walking path using this model.

### 3.2. Abathic distortion

Another possible cause for systematic errors is that the observer builds a distorted 3D representation of space when they are at the start zone and uses it to guide their pointing. Many models of binocular space perception assume that there is a distorted mapping between true space and represented space, often with an ‘abathic’ distance at which objects are judged to be at the correct distance and a compression of visual space towards this plane [44, 45] as illustrated in Fig. 5a. Applying this 2-parameter model to our pointing data, the best-fitting abathic distance is actually about 5 m *behind* the observer when they view the scene (i.e. behind the start zone, Section S4.3). Another problem for this model, just like the noisy-path-integration model, is that it makes no prediction about the effect of the slant of the obscuring wall.

### 3.3. Retrofit

The abathic distance model is only one case of a model in which the observer uses a distorted 3D model of the scene as the basis of their pointing responses. An extreme version of this type of model is to allow the location of the target boxes to vary freely in order to find a scene layout that maximizes the likelihood of the pointing data, i.e. ‘retrofitting’ the scene to match the data. As such, it is hardly fair to describe this a ‘model’ in the sense of having any predictive value since the 3D configuration of the supposed internal representation is determined entirely by the data. The result of doing this for *Experiment 1* is shown in Figs. 6a–6d. We used the pointing directions of all 20 participants from 3 pointing zones per target box in each condition to calculate the most likely location of the box that would explain their pointing responses. Fig. 6a shows the result of this calculation for the blue box in one condition. It is clear that the derived box location is far from the true location. Figs. 6b, 6c, and 6d show the same for the pink, red and yellow target boxes respectively (see Section S4.4 for details and Fig. S5c which shows the same retrofit process applied to the results of *Experiment 3*). Fig. 6e marks the location of the peaks of the likelihood distributions illustrated in Figs. 6a–6d and all other conditions in *Experiment 1*. These points tend to cluster around a plane that is parallel to the plane of the obscuring wall whereas the true locations of the target boxes (shown in translucent colors in Fig. 6f) are nothing like this. As mentioned earlier, participants tend to point in a direction that is more orthogonal to the obscuring wall than the true target direction, but now it is clear that the inferred target locations all have a similar depth relative to the obscuring walls, as if the depth structure of the scene has been ‘squashed’. In reality, the pink and blue boxes were very close to the obscuring wall in the center of the room, while red and yellow boxes were more distant. Yet, according to participants’ pointing data, the *apparent* location of the targets were all at a similar distance relative to the wall.

**Figure 6:**
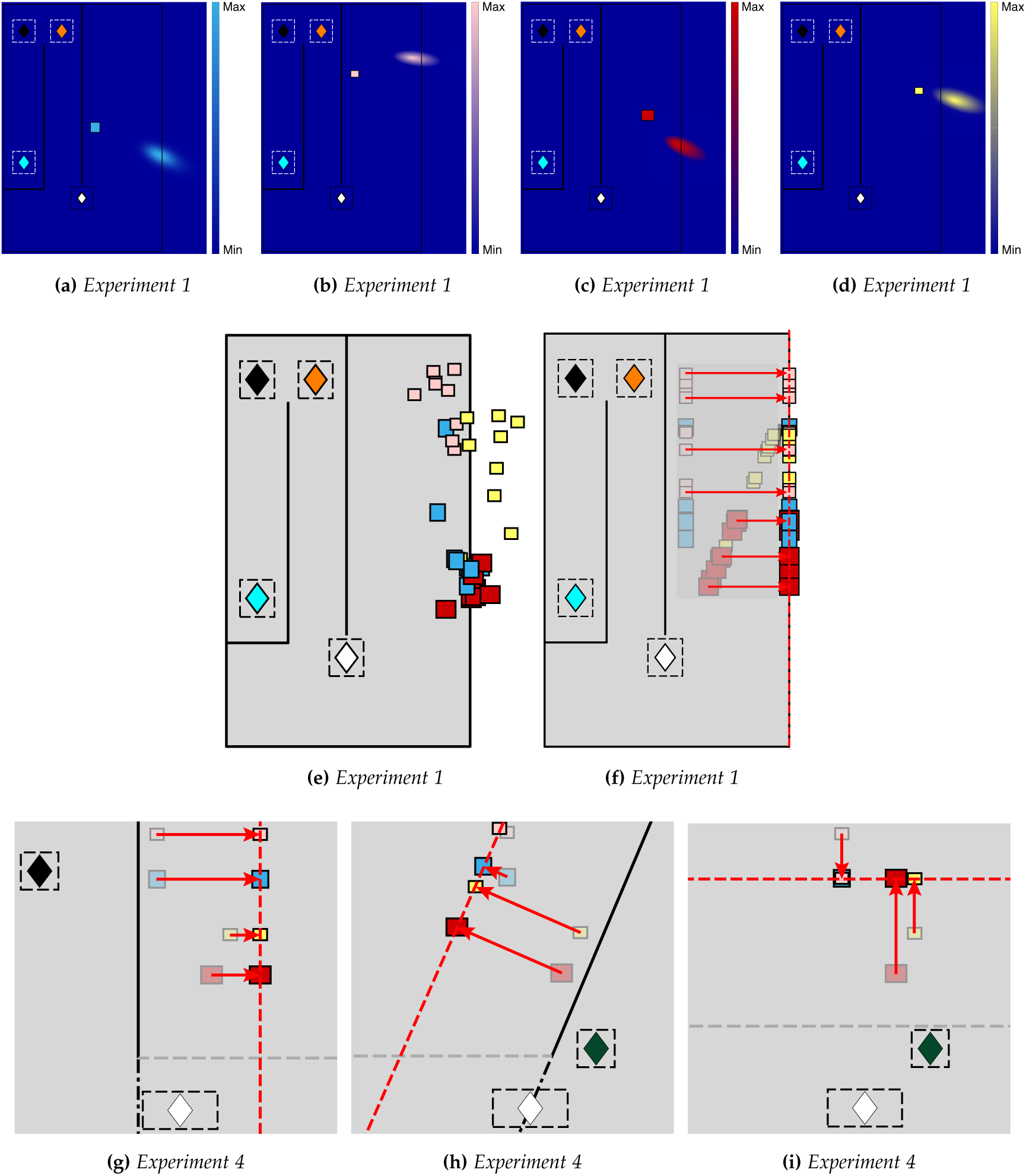
Retrofit target locations and the inspiration for the projection plane ‘model’. (a–d) Using all the pointing directions of all participants from three pointing zones it is possible to compute the most likely location of a target that would generate this pointing directions. Likelihood is plotted at every location and its peak is visible at the maximum-likelihood location. (e) Plan view of the maximum-likelihood positions of all boxes in each layout. Most of these locations are shifted towards a plane that is parallel to the central obscuring wall. (f) The projection plane model assumes that the projection plane is parallel to the wall, with a set distance (1.77 m), derived from the data in Experiment 1. (g–i) Illustrations of the projection-plane model applied to different configurations in Experiment 4.

### 3.4. Projection plane

So, as a *post-hoc* ‘model’ of this apparent distortion of the scene structure, we can apply the following prediction to the other experiments. Participants will point to remembered targets by assuming that all the targets in that experiment lie on a single plane that is parallel to and 1.77 m behind the obscuring wall. The value of 1.77 m is simply taken from the best fit to the points shown in Fig. 6e. Fig. 6f, Fig. 6 g, Fig. 6 h and Fig. 6i show examples of this projection-plane rule applied in other conditions.

### 3.5. Model comparison

We have already discussed the fact that the noisy-path-integration and abathic-distance-distortion models cannot account for the effect of the obscuring wall orientation (Fig. 4). Table S1 shows the residuals for these two models and for the projection-plane model for Experiment 1, where all three models are fitted to the data of all participants combined. The residuals are lower for the projection-plane model for 17/20 participants but the key test for this model is to see whether it can predict performance in *Experiments 2* to *4* where it provides a zero-parameter prediction of the pointing data.

The retrofit model is rather different. It is provided with all the data and asked to derive a 3D structure of the scene that would best account for participants’ pointing directions using a large number of free parameters (24) so, unlike the projection-plane model, it does not make predictions. Table S2 shows a measure of goodness of fit of the projection plane model and the retrofit models, adjusted for the number of parameters in each (using Bayesian (BIC) and Akaike (AIC) Information Criteria). According to the BIC, the projection plane model outperforms the retrofit method in all 4 experiments. This can be seen graphically in Fig. 7, which shows the predicted pointing errors from the retrofit model plotted against the actual pointing errors, while Fig. 8 shows the same for the projection-plane model: the slopes are all steeper for the projection plane model.

**Figure 7:**
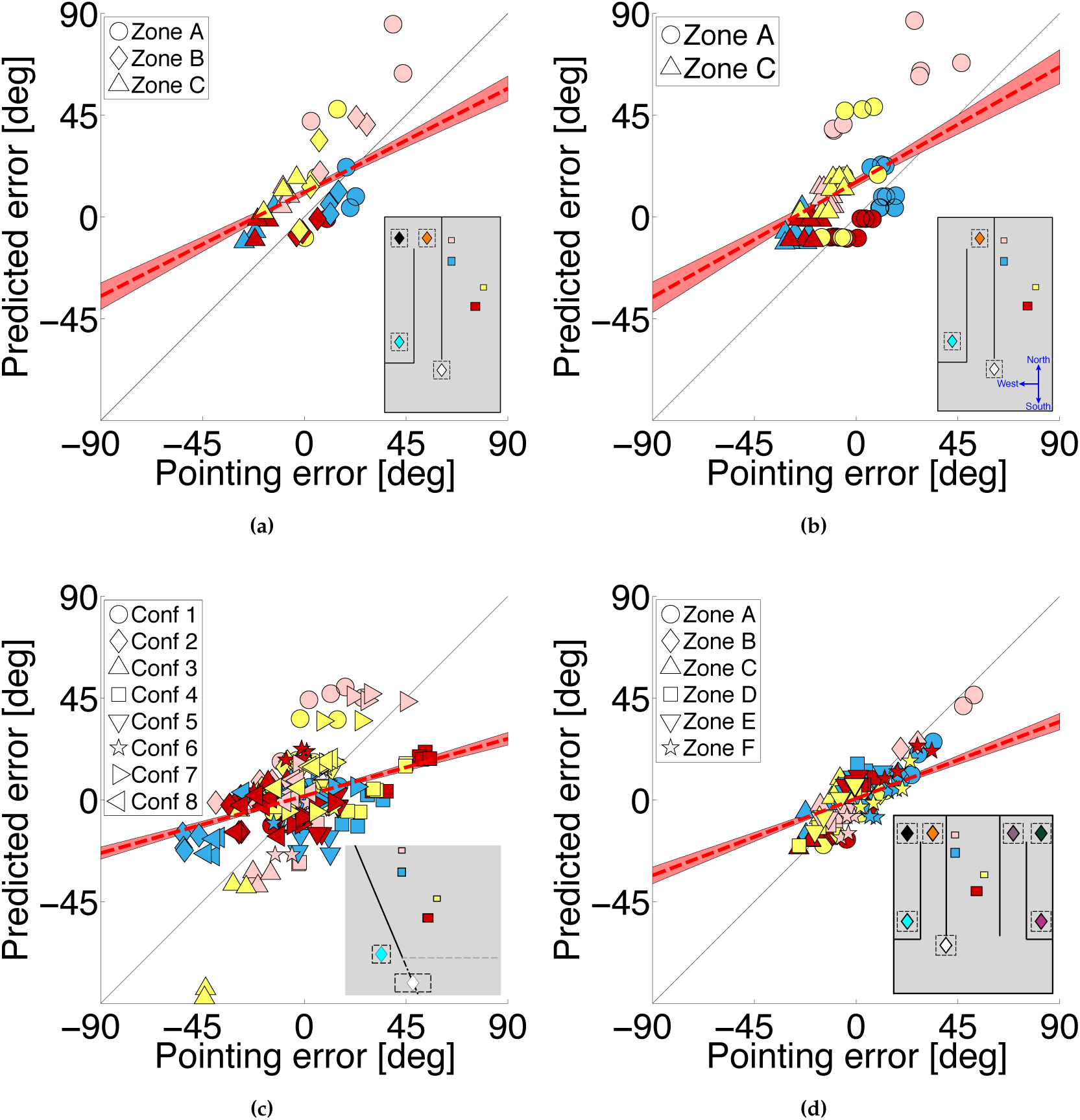
Predictions of the ‘retrofit’ model. For box layouts that were shown in Experiments 1, 2 and 4, a configuration of box locations was determined that would best explain the pointing data (see Fig. S5b). Predictions of the model are shown for (a) Experiment 1, (R = 0.52, p < 0.001, RMSE = 24.1°, slope = 0.51), (b) Experiment 2, (R = 0.54, p < 0.001, RMSE = 29.3°, slope = 0.57), and (c) Experiment 4 (R = 0.32 and p = 0.0014, RMSE = 34.1°, slope = 0.28). Scene configurations are shown in Fig. S1, (d) Experiment 3 was treated separately as the box layouts were different (see Fig. S5c) and all 6 pointing zones were used to predict box locations (R = 0.40 and p < 0.001, RMSE = 25.7°, slope = 0.38).

**Figure 8:**
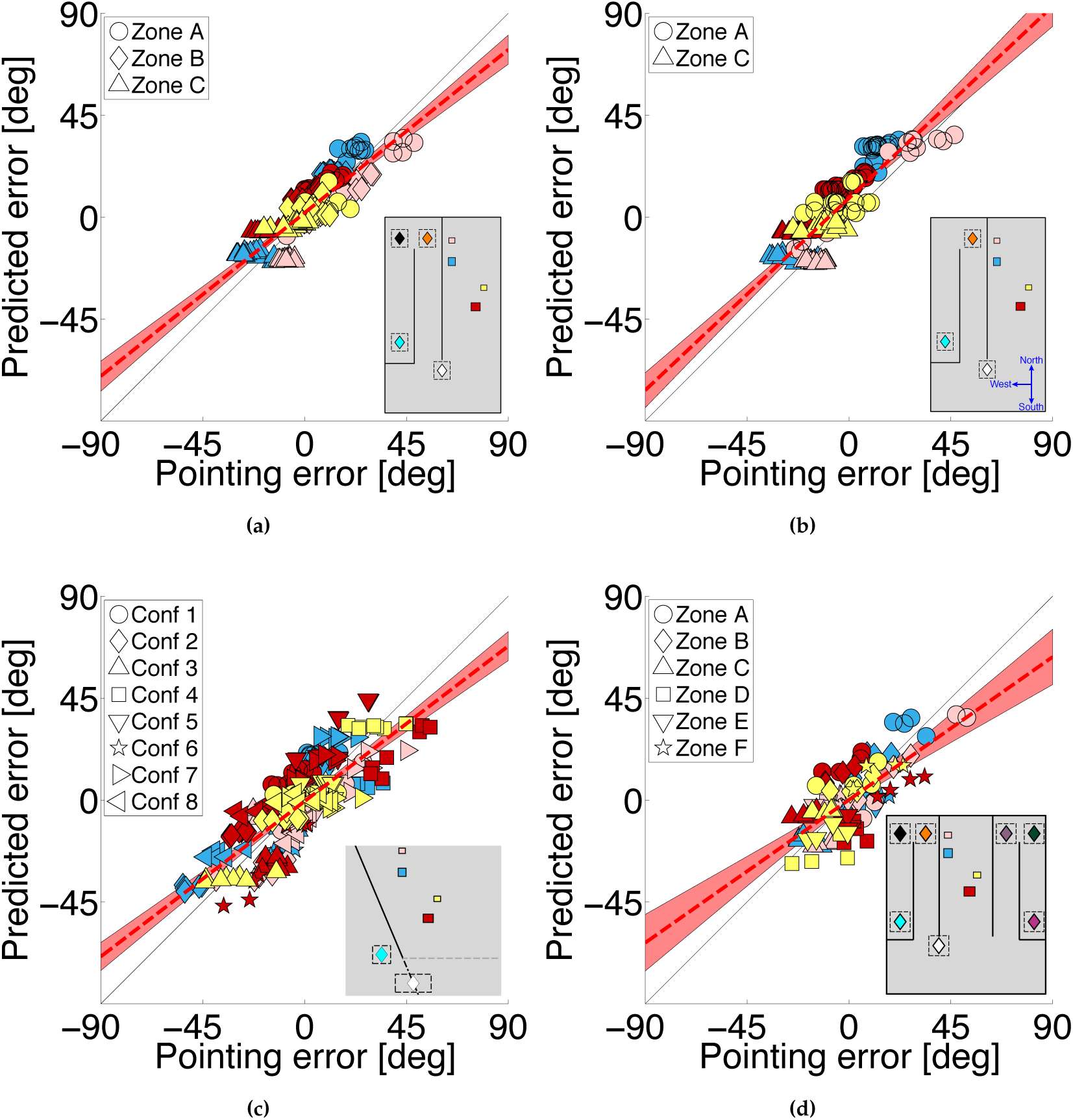
Predictions of the projection-plane model. As for Fig. 7 but now showing the pointing errors predicted by the projection-plane model. (a) Predictions for Experiment 1 (R = 0.86 and p < 0.001, RMSE = 8.6°, slope = 0.80), (b) Experiment 2 (R = 0.86 and p < 0.001, RMSE = 12.2°, slope = 0.89), (c) Experiment 4, (R = 0.73 and p < 0.001, RMSE = 12.2°, slope = 0.73) and (d) Experiment 3 (R = 0.81 and p < 0.001, RMSE = 11.8°, slope = 0.66)

Finally, returning to Fig. 4, Figs. 4j and 4i show that the projection-plane model accounts for the bias that is caused by the ‘slanted’ versus the ‘East-West’ wall in that experiment. Using paired trials (i.e. with everything else kept the same except for the wall orientation), these plots show the distribution of differences in pointing directions relative to the *predicted* direction of the target. Unlike the original biased pointing directions (Figs. 4 g and 4f), the mean of the distribution is no longer significantly different from zero (*M* = −1.81, *SD* = 22.0, *t*(319) = −1.48, *p* = 0.140), showing that the model has accounted well for the pointing biases.

## 4. Discussion

For a spatial representation to be generated in one location and used in another, there must be some transformation of the representation, or its read-out, that takes account of the observer’s movement. The experiments described here show that in humans, this process is highly inaccurate and the biases are remarkably consistent across participants and across many different conditions.

The data appear to rule out several important hypotheses. Crucially, any hypothesis that seeks to explain the errors in terms of a distorted internal model of the scene at the initial encoding stage will fail to capture the marked effects of the obscuring wall, which is generally not visible at the encoding stage (see Fig. 4). We examined standard models of a distorted visual world, namely compression of visual space around an ‘abathic’ distance [44, 45] (details in Section S4.3) but even when we allow *any* type of distortion of the scene that the observer sees from the starting zone to explain the data (provided that the *same* distortion is used to explain a participant’s pointing direction from all pointing zones and any wall orientation), this type of model still provides a worse fit to the data than our projection-plane model (see Fig. 7, 8 and Table S2). The post-hoc nature of the ‘retrofit’ model could be considered absurdly generous. The fact that it is *still* worse than a zero-parameter model generated from the first experiment provides strong evidence against 3D reconstruction models of this type.

We also examined and ruled out a hypothesis based on noisy path integration [46] as an explanation of our data. One strong piece of evidence in relation to such hypotheses is the high correlation between the magnitude of errors participants make when they walk to a pointing location either via a long or a short route (Fig. 2d and Fig. S2a). These data suggest that people point to a memory of the target boxes that is quite different from their true locations but it is not the route to the pointing zone that can explain this. Instead, it is something about the location and the scene when the participant gets there.

Biases in pointing to unseen targets have also been shown to be dependent on gaze direction [47]. Participants in our experiment tended to look in the direction to which they were pointing and it seems unlikely that one could predict the pattern of biases we observed on the basis of a systematic difference between gaze direction for the different target boxes. The strong effect of wall orientation and the absence of effect of body orientation militate against this suggestion.

One paper that uses a similar paradigm to ours also shows consistent pointing biases [26]. When the pointing zone (in their experiment, called an ‘indirect waypoint’) was displaced from the location at which the scene layout was learned, pointing was shifted consistently in the direction of the waypoint. The participant was blindfolded during the test phase and guided to the waypoint so there was no equivalent of our wall and only two waypoints were tested so it is hard to make a direct comparison with our data. Nevertheless, their data suggest that, in line with our data, the location of the pointing zone with respect to the start zone has a systematic biasing effect on pointing directions. It has been suggested that ‘categorical bias’ could account for our data, where participants point somewhere between the true location of a target and the centre of a previously-defined region [48, 49]. If the category was the centre of all the box locations (or all the box locations of one colour) this would not give rise to the pattern of biases we observed. If ‘categories’ are supposed to be built up across pointing zones then a strong prediction of this model is that repeating the experiment with only one pointing zone would abolish the biases, a result we believe to be unlikely.

It has often been shown that the observer’s orientation can have a significant effect on pointing performance. For example, when the orientation of the participant in the test phase differs from their orientation during learning of the scene layout this can influence pointing directions (for both real walking [50] or imaginary movement [17, 51] in the test phase). Meilinger *et al*. [52] investigated the effect of adding walls to an environment and showed they have a significant effect. However, the authors did not examine the biasing effects of moving to different pointing locations nor can the results be compared directly to ours since they report ‘absolute errors’ in pointing, a measure that conflates variable errors and systematic biases. In fact, this is a common problem in the literature; many other papers report only absolute errors in pointing [53, 24, 17, 21, 54, 22, 55, 19]. Röhrich *et al*. [32] showed that the participant’s location when pointing was likely to be important in determining the reference frame that they used. In this experiment, participants pointed to a well-known market square from different locations in the town. Pointing location affected the orientation in which participants drew a plan-view sketch of the square. However, the authors did not predict biases in pointing. There are also claims that egocentric factors play a role in pointing biases [17, 39] but we did not find any effect in our experiment: pointing errors were unaffected by a 180° change in the observer relative to the scene at the moment they discovered which target they should point at (Fig. 3b).

‘Boundaries’, such as the walls in our experiment, have often been considered as important in determining the coding of spatial positions [56, 57, 58] although these earlier papers do not provide predictions about the bias in representation of objects the other side of wall. Opaque boundaries can have an effect on the way that hidden objects are categorised. For example, spatial memories can be biased towards the centres of enclosed regions [59] (a type of categorical bias, as discussed above) but there are no enclosed regions in our experiment and so it is hard to see how this could explain the biases we report here. We include all the pointing data online (Section S6) with code to produce Figures 7 and 8 so testing alternative models should be relatively straightforward.

The fact that the most lenient 3D reconstruction model does not do well in explaining our data raises the question of what participants do instead. The ‘projection-plane’ model is essentially a re-description of the data in one experiment, rather than an explanation, albeit one that then extends remarkably successfully to other situations (‘remarkably’ because it is a zero-free-parameter model). If one were to come up with an image-based model that gave rise to the same pattern of biases as participants, it seems that it would have to start with an image of the scene as viewed by someone facing the obscuring wall (similar to studies of people viewing pictures obliquely [60]), even though the participant never sees the scene from this perspective. Supposing, for the moment, that they could construct such an image ‘painted’ on the obscuring wall (an important and problematic caveat) then, as the participant walks from, say, pointing zone B to zone C they would generate perspective distortions of this ‘image’ which would be, of course, very similar to the image distortion of the ‘projected plane’ described in our model. Adding a fixed rotation of the pointing direction - the same for all targets - so that the pointing directions are all to objects behind the wall, would then lead to a set of pointing directions that are very similar to the ‘projection plane’ model we have described, even though it is not based on a 3D model.

If the visual system does use some heuristic like this rather than generating a 3D model of the scene then, whatever heuristic it uses, it seems to ignore critical aspects of the geometry of the scene, such as relative depths. The consequence is that objects are treated in a more simplistic way (equivalent to assuming that the target objects all lie on a plane) than would be the case if the internal model behaved according to the geometry of a real 3D model. There is increasing interest in models of processing at all levels, from a ‘2½D sketch’ [61, 62] through to pointing and navigation, that avoid the necessity for building a 3D model of surfaces or scene layout [63, 64]. Recent findings using ‘Generative Query Networks’ [8] suggest that we may see rapid advances in our understanding of the process of ‘imagining’ novel views of an observed scene and how this might be achieved without using a 3D reconstruction.

Finally, it is worth considering what effect such large biases might have in ordinary life. The most relevant data from our experiments in relation to this question are, arguably, the pointing biases that we recorded from the very first trial for each participant. We made sure that, on this first trial, the participant was unaware of the task they were about to be asked to do. Biases in this case were even larger than for the rest of the data (Fig. S2 h). If these data reflect performance in daily living, one might ask why we so rarely encounter catastrophic consequences. However, the task that we asked our participants to carry out is an unusual one and, under most circumstances, it is likely that visual landmarks will help to refine direction judgments *en route* to a target.

## Conclusion

Our conclusions are twofold. First, although human observers *can* point to remembered objects, and hence must update some form of internal representation while the objects are out of view, we have shown that they make highly repeatable errors when doing so. Second, the best explanation of our data is not consistent with a single stable 3D model (even a distorted one) of the target locations. This means that whatever the rules are for spatial updating in human observers, they must involve more than the structure of the remembered scene and geometric integration (even with errors) of the path taken by the observer.

## Acknowledgements

This research was funded by a *Microsoft Research* PhD Scholarship to JV and by *EPSRC* grant EP/K011766/1 and *Dstl/EPSRC* grant EP/N019423/1 to AG. We thank Bruce Cumming for discussions that led to Experiment 4 and Lyndsey Pickup for help with modeling.

## Author contributions

Conceived project: A.G. and A.W.F.; contributed ideas to experiment: A.G., J.V. and A.W.F.; development of testing platform and environments: J.V.; performed experiments and analysis: J.V.; contributed ideas to modeling: A.W.F., A.G. and J.V.; wrote paper: A.G. and J.V.

## Supplementary Material

### S1. General Methods

In virtual reality, participants viewed the stimuli through a binocular head-mounted display NVIS SX111 with horizontal view of 102°, vertical view of 64° and binocular overlap of 50°. Each display for the right and the left eye had a resolution of 1280×1024 pixels refreshed at a rate of 60 Hz and an end-to-end latency of 2 frames. The tracking system consisted of 14 MX3/T20/T20-S Vicon tracking cameras, a tracking computer (Quad Core Intel Xeon 3.6 GHz processor, NVidia Quadro K2000 graphics card, 8GB RAM) running Windows and a graphics computer (8 core AMD Opteron 6212 processor, dual NVidia GeForce GTX 590 graphics cards, 16GB RAM) running Linux that generated the experimental stimulus. The head-mounted display was tracked at a rate of 240 Hz.

In the real environment, participants wore a tracked hardhat with blinkers on the side to imitate the horizontal field of vision of the head-mounted display. The participants were blindfolded when guided to the start zone by the experimenter. The target boxes had markers stuck to them so that they could be tracked and their position recorded during trials. In both virtual and real environments, participants used a hand-held, fully-tracked pointer resembling a gun, to indicate a pointing direction with the direction of the pointer and with a button press.

All participants had normal or corrected-to-normal vision and could distinguish the colors of the target boxes. Their acuity and stereoacuity were tested before the start of the experiment: all had 6/6 vision or better and normal stereopsis with 60 arc seconds or better.

### S2. Experiments

#### S2.1. Ethical Approval

The experiments described in this study received the approval of the University of Reading Research Ethics Committee. Informed consent was obtained from all participants.

#### S2.2. Experiment 1

There were two parts to the experiment. In one, carried out in virtual reality, on different trials participants walked via two different routes (‘direct’ and ‘indirect’) to get to the pointing zones from which they pointed to the remembered targets. In the other, participants carried out one of these tasks (‘indirect’ route) in a very similar environment in the real world. The very first trial the participants experienced was one in the real world.

##### Participants

For the first experiment, we recruited 22 participants (aged 19–46) who were paid £10.00 per hour. Data from 2 participants had to be discarded as one failed to understand the task and one felt too uncomfortable wearing the head-mounted display. 19 out of the 20 were naïve to the experiment, and 1 was a researcher in our lab who had prior knowledge about the task (see Table S1).

##### Procedure

In virtual reality, in order to start a trial, participants walked towards a large green cube with an arrow pointing towards it in an otherwise black space then, as soon as they stepped inside the cube, the stimulus appeared. In the real world, participants wore a blindfold and were guided to the start zone by the experimenter. As soon as they were in the center of the start zone, the experimenter removed the blindfold. In both real and virtual experiments, the participants then viewed 4 boxes from this start zone. They were told that from where they were standing, the blue box was always closer than the pink box and both were in the same visual direction. Likewise, the red box was always closer than the yellow box and, again, both were in the same visual direction when viewed from the start zone (see the example plan view in Fig. 1a–c). Participants took as much time as they needed to view and memorize the box positions, although this was typically between 10 and 20 seconds. If they attempted to leave the start zone to walk closer to the target boxes, the whole scene disappeared in virtual reality or were inhibited by the experimenter in the real world. When they had finished memorizing the box layout, they walked behind a ‘wall’ (partition) towards a pointing zone. The layout of the partitions is shown in Fig. 1. The ‘inner’ north-south partition, shown by the dashed line in Fig. 1c, was removed in the ‘direct’ walking condition (only tested in virtual reality). The indirect and direct walking conditions were intermingled and tested in a randomised order. The participants did not know which condition they were being tested in at the beginning of each trial while standing at the start zone. In virtual reality, the pointing zone was indicated by a colored poster (colored according to the color of the box to which they should point) and this poster only appeared after they left the start zone. Similarly, in the real-world task, there were three white posters at the three pointing zones (see Fig. 1c) and the participant was only told which one to stop at after they had left the start zone. In virtual reality, the poster changed its color after each shot to indicate the next target box to point at. In the real world, the experimenter told the participant which box to point at. Following these instructions, the participant had to point 8 times to each of the 4 boxes in a pseudo-random order, i.e. 32 pointing directions in all. The end of each trial was indicated by the whole scene disappearing in virtual reality (replaced by the large green box) and in the real world the experimenter told the participant to wait to be guided back blindfolded to the start. The participant received a score ranging from 0–100% reflecting the accuracy of all the shots as this increased participants’ motivation) but this information could not be used to infer the direction or magnitude of their pointing error on any given shot.

##### Box layouts and stimulus 8

participants were tested on 9 box layouts (1,440 pointing directions in total in VR per participant: ‘Indirect’ — 9 layouts × 4 boxes × 3 zones × 8 shots, ‘Direct’ — 9 layouts × 4 boxes × 2 zones × 8 shots), whereas the remaining 12 participants were tested on 4 layouts (640 trials in VR per participant). All participants repeated the ‘indirect’ condition in the real world in exactly the same way as they did in VR. All the box layouts are shown online in an interactive website, see Section S6, including the raw pointing data in each case. The virtual and the real stimulus were designed to be as similar as possible, with a similar scale, texture mapping taken from photographs in the laboratory and target boxes using the same colors and icons, see Fig. 1a–b.

The box positions in the 9 box layouts were chosen such that the following criteria were satisfied, see Fig. 1(c):

- The blue and the pink boxes were positioned along one line (as seen in plan view) while the red and the yellow box were positioned along a second line. The two lines intersected inside the start zone.
- All box layouts preserved the box order: this meant that the blue box was always in front of the pink box, and the red box was in front of the yellow box, but the distances to each box varied from layout to layout.
- The 2 lines were 25° apart from one another. This number allowed a range of target distances along these line while the boxes all remained within the real room.

#### S2.3. Experiment 2

This experiment aimed to identify the influence of the facing orientation at the pointing zones. 7 participants, all of whom had been participants in the first experiment, viewed the same 6 layouts as in *Experiment 1* but in virtual reality only. They then walked to either zone A or zone C, stopped and faced a poster located either to the ‘North’, ‘South’, or ‘West’ of the pointing zone (unlike *Experiment 1* when there was only one poster position per pointing zone, see Fig. S1a. Otherwise, the procedure was identical to *Experiment 1*, with the exception that the participant pointed 6 times (rather than 8) to each of the boxes in a random order at the pointing zones. Data are shown in Fig. S2 and Fig. 3b.

#### S2.4. Experiment 3

The third experiment was also carried out only in virtual reality and here the testing room was scaled (in the x-y plane) by a factor of 1.5, i.e. the height of the room did not change. The pointing zones A, B and C were mirrored along the center line of the room to create an additional 3 pointing zones D, E and F, see Fig. S1b but the physical size of the laboratory dictated that participants could only go to one side of the scene or the other (either zones A, B, C or zones D, E, F). The participants did not know whether they would walk to the left or to the right from the start zone until the moment that they pressed the button (when the boxes disappeared). There were 6 participants, 3 of whom had taken part in the previous experiments while 3 were naïve.

#### S2.5. Experiment 4

The scene layout for this experiment is shown in Fig. S1c–j. There were 4 pointing zones B, C, D and E and only 4 box layouts. All the data were collected in virtual reality. For every condition (box layout and pointing zone), participants carried out 2 trials with the wall in one orientation and the same number of trials with the wall in a different orientation (i.e. 6 shots per target for each of the targets, layouts, zones and repeated for 2 runs).

4 naïve and 6 non-naïve participants were recruited for this study. The procedure of this experiment was similar to *Experiments 1–3* with the following differences: Participants were standing at the start zone and viewed the target box positions. After pressing a button, a 1.8 m tall wall appeared in front of them, see gray dotted line in Fig. S1c–j. A white poster at eye height and a green square on the floor appeared in the room indicating the location of the pointing zone and the orientation. When the participant left the start zone, another wall appeared, see black solid line in Fig. S1c–j. The rest of the trial was as described in *Experiment 1–3*.

**Figure S1:**
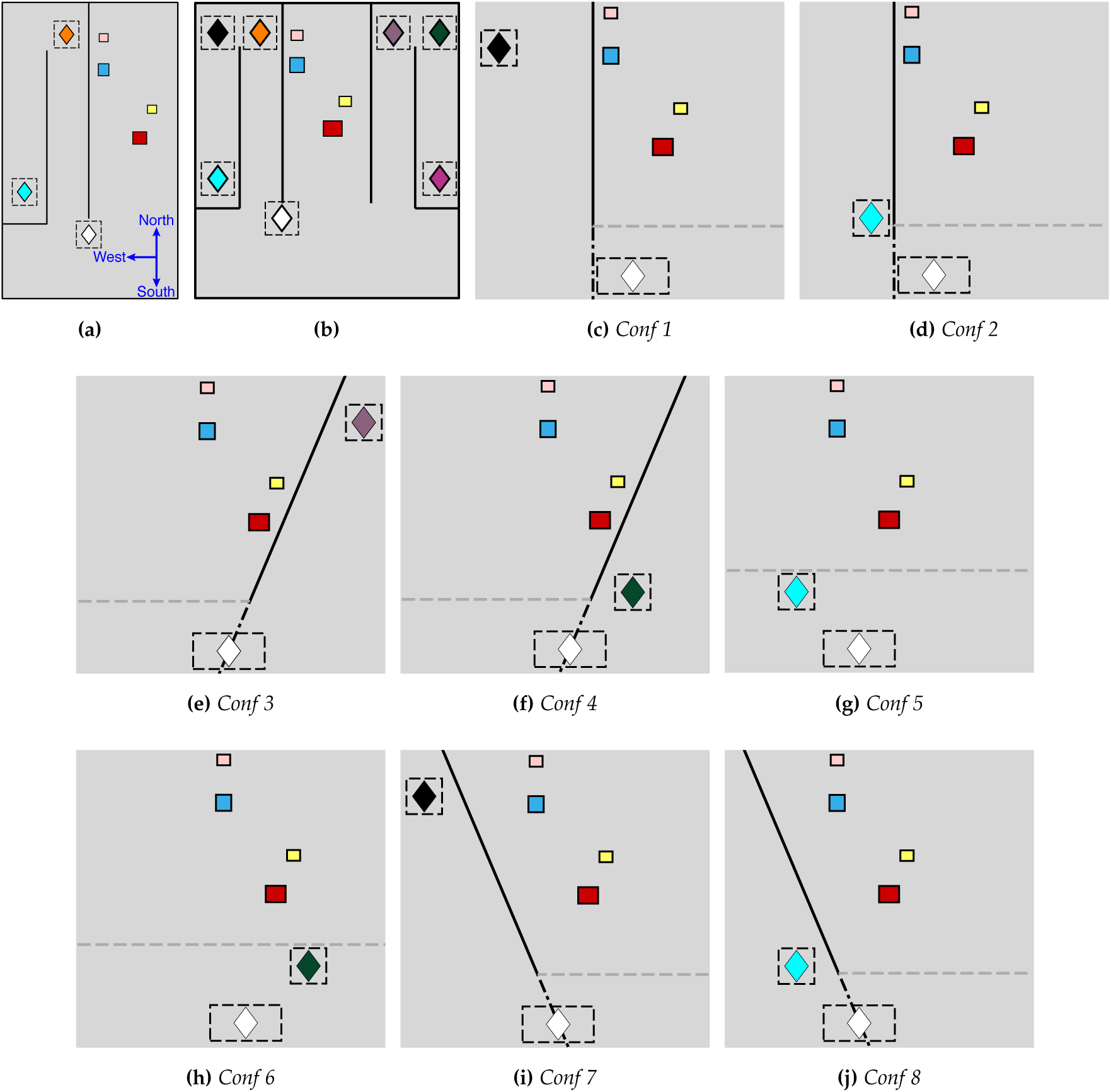
Plan view of (a) Experiment 2, (b), Experiment 3 and (c–j) Experiment 4 with 8 different configurations. Black solid line indicates walls that were always present. In (c–j), the gray dotted line shows the wall that appeared when the participant pressed a button at the start zone (white diamond). The black dotted line shows the wall that appeared after participant left the start zone. In (g) and (h) the wall remained in the same place.

### S3. Results

#### S3.1. Individual participant data

**Figure S2:**
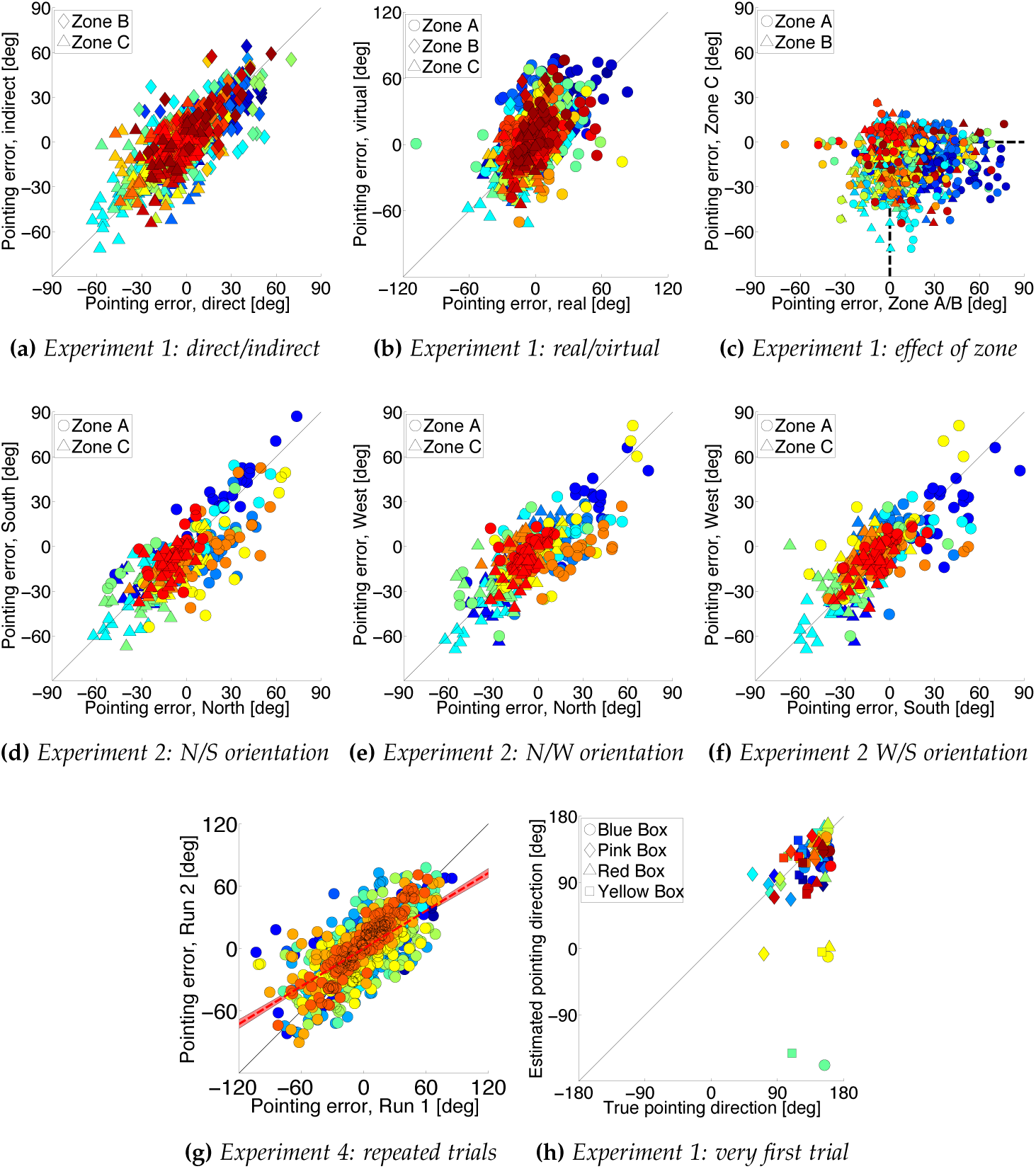
Individual participant data. Colors indicate different participants. (a) corresponds to Fig. 2 (direct versus indirect walking), (b) to Fig. 3a (real versus VR), (c) corresponds to Fig. 3c (zone C versus zone A/B) and (d) corresponds to Fig. 3b (‘North’-facing versus ‘South’-facing). (e) and (f) show data from an additional condition (participants initially faced ‘West’) compared to the ‘North’ and ‘South’ conditions. (g) In Experiment 4, the same conditions were repeated twice; pointing errors for the two runs are plotted against each other (slope is 0.61, indicating a smaller range of errors on the second run). (f) The very first time that participants viewed the stimulus in the real world, they were given no instructions at the start zone. Then, at the pointing zone, they were asked to point to the four boxes in a random order, eight times each (i.e. 32 shots per participant). It is debatable whether a post-hoc power analysis is of value in this instance but, for the record, this shows that the power achieved to rule out the correlation shown in Fig. 2d/FigureS2a occurring by chance is, to a very close approximation, 100%. More relevantly, the same pattern of biases is found throughout the remaining experiments in the paper.

#### S3.2. Alternative definitions of pointing direction

**Figure S3:**
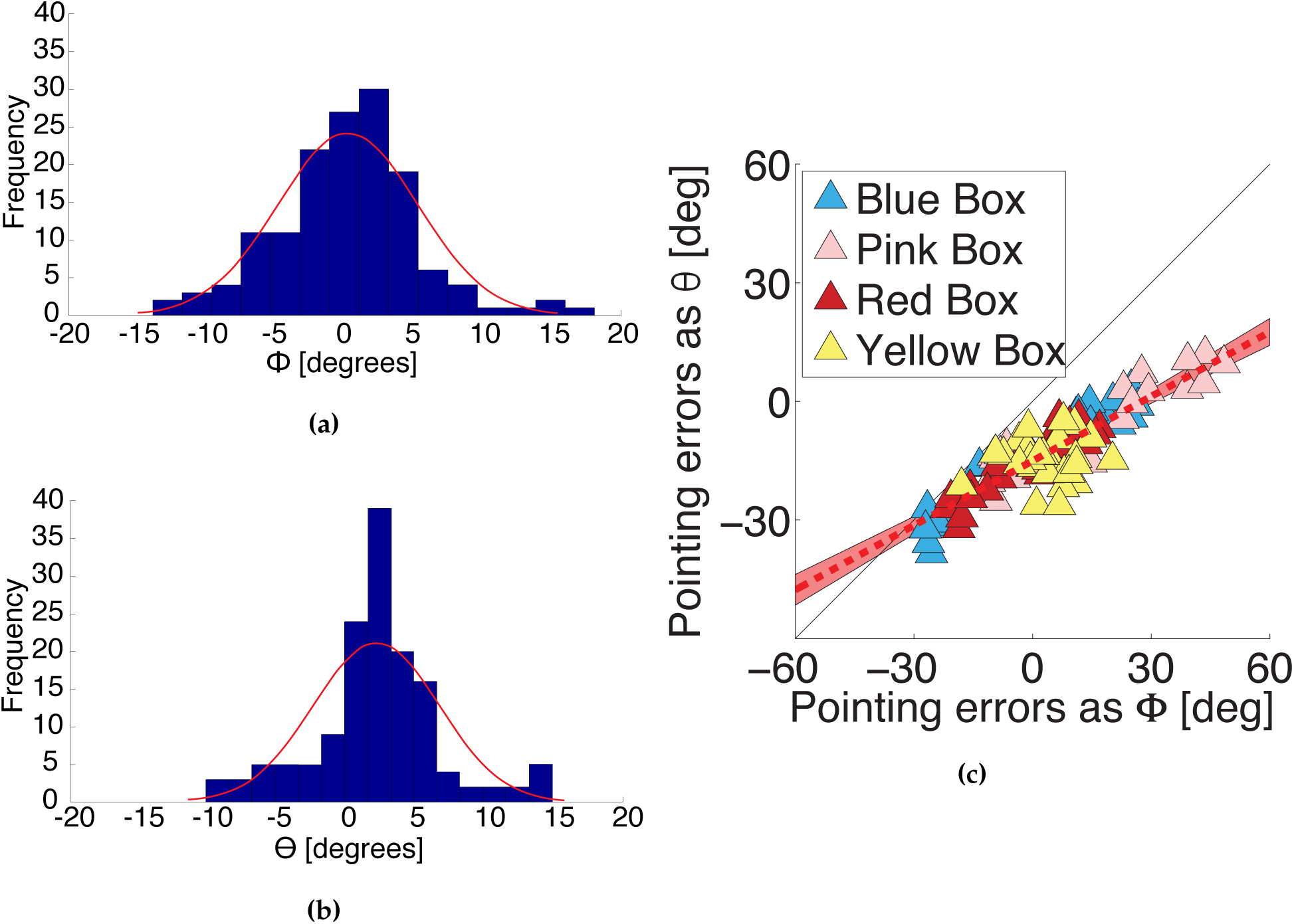
Evidence supporting the choice of ‘shooting’ angle (ϕ) rather than visual direction (θ) as the most appropriate definition of pointing direction. In a control condition, participants pointed at target boxes from the start zone so the target was visible (18 participants tested on 2 box layouts with 4 target boxes in each layout, 144 shots in total). Their instruction was to ‘shoot at the box’, just as it was in the spatial updating experiments. (a) Distribution of pointing errors when target direction is defined relative to the pointing device and shooting direction is defined by the orientation of the device (see ϕ in Fig. 1d). (b) As for (a) but with pointing errors defined as the difference in visual direction of the target and the pointing device as measured from the cyclopean point (see θ in Fig. 1e). The mean of the distribution is significantly biased for θ (t-test, p < 0.001) but not for ϕ (p = 0.605) suggesting ϕ reflects participants’ intentions when pointing. (c) There is a significant correlation between the ϕ and θ measures in the spatial updating experiments (correlation coefficient 0.91). These data are from Experiment 1. When ϕ = 0, θ is about −20°. A negative bias is the direction of bias that would be created if a right-handed participant held the pointing device slightly out to their right and pointed to a target directly ahead of them.

### S4. Visual Models

#### S4.1. Noisy-path-integration model

The noisy-path-integration model simulates a moving observer storing an egocentric map of the box positions by constantly updating the heading direction with respect to ‘North’ (*α*) and estimating the distance traveled on each step (*d*). Here, we assume that the observer misestimates *α* and *d* with a constant error (multiplicative calibration errors *ω_α_* and *ω_d_* respectively). This leads to a cumulative error in the estimate of the participant’s location. The box locations are assumed to be known correctly.

Initially, the position of the boxes is given, by definition, as follows, where 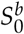 is the starting distance of box *b* and angle 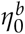 is its visual direction with respect to ‘North’ 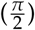 as viewed from the start position (boxes are still visible here):

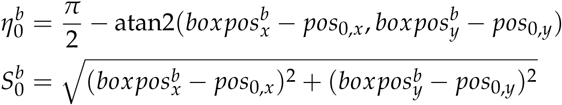

with 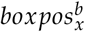 and 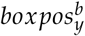 being the *x*- and *y*-coordinates of the *b^th^* box, *pos*_0,*x*_ and *pos*_0,*y*_ the cyclopean point of the observer at the start position. At subsequent steps, when the boxes become obscured by the wall, the polar coordinates are calculated using the following equations:

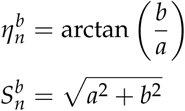

with

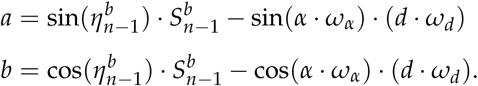

We fitted this model to all the data from all participants in Experiment 1 by varying the two free parameters, *ω_α_* and *ω_d_*, to give the minimum root-mean-square-error between the actual and predicted pointing directions. (see Fig. S4a).

**Figure S4:**
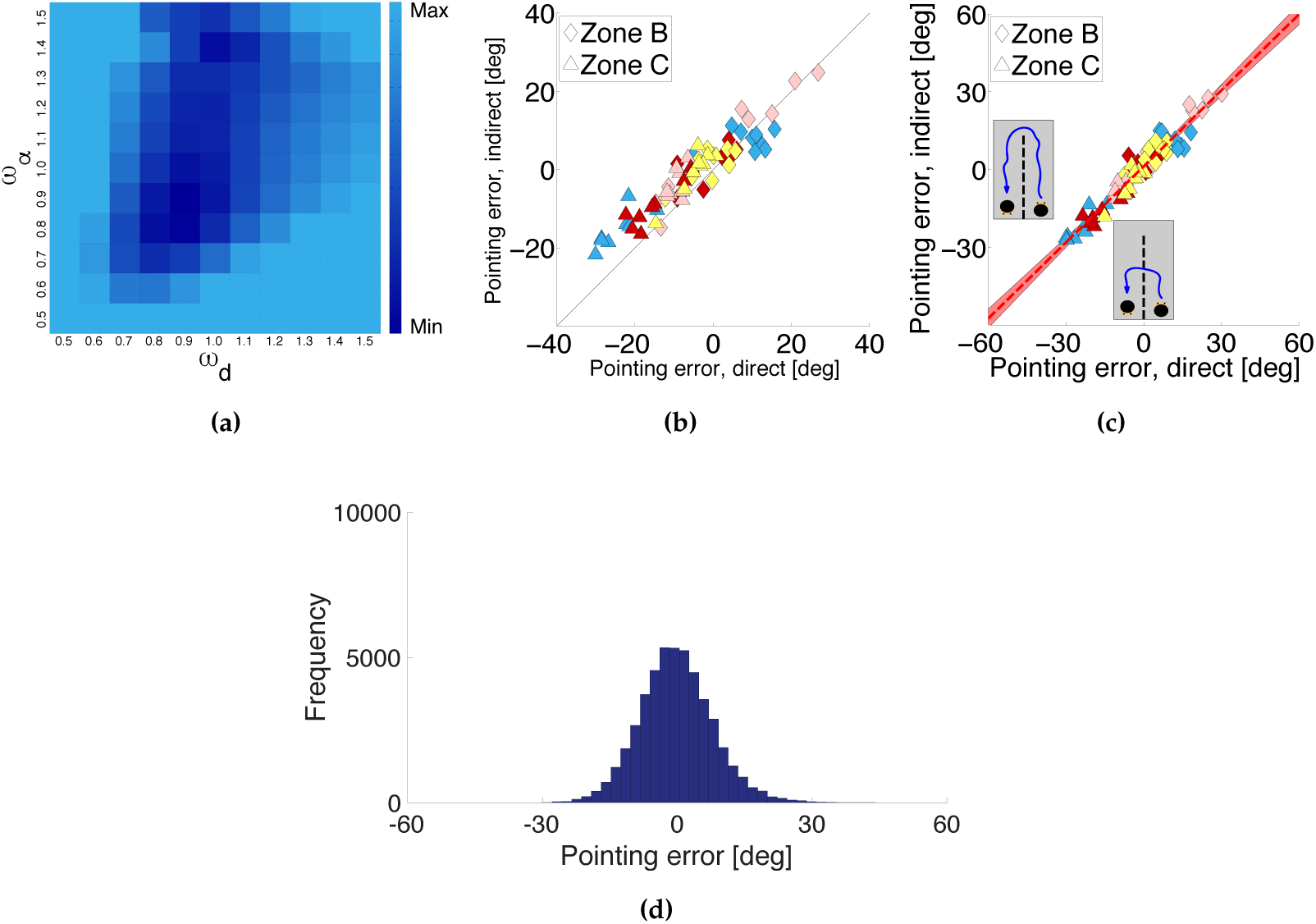
Illustration of the noisy-path-integration model. (a) Using the data from all participants, this plots shows the different values of the root-mean-square error (RMSE) between data and model measured in degrees using different combinations of the model parameters ω_d_ (distance bias) and ω_α_ (angular bias). Any combination with an RMSE larger than twice that of the ground truth model for the indirect condition of Experiment 1 (RMSE_lim_ = 2 · 16.6°) has been set to that value (‘Max’). Dark blue squares show the calibration errors with lower RMSE values, the lowest is at (0.9, 0.9) with an RMSE value of 13.2° (‘Min’). (b) Comparison of noisy-path-integration predictions for direct and indirect walking. Error predicted by the noisy-path-integration model for the indirect walking condition plotted against this value for direct walking trails. As in Fig. 2d, each symbol is based on the mean data for 20 participants. The data for zone C (triangles) is most informative as the length difference between the direct and indirect paths is most extreme in this case. Here, the errors for indirect walking are significantly more positive than the errors for direct walking (direct walking M = 6.06, SD = 3.19, indirect walking M = 0.764, SD = 2.76, t(35) = 18.5, p < 0.001), whereas the experimental data for these two conditions, reproduced here in (c) from Fig. 2d, were not significantly different. (d) Histogram of prediction errors calculated 100 times for each box in each layout, using the walking trajectory of every participant tested in the indirect walking condition of Experiment 1 at one of the pointing zones (zone C) with a normally distributed random noise on the estimate of 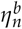. The mean of the pointing errors is not significantly different from zero. The mean of the distribution for zone A and zone B was also not significantly different from zero.

#### S4.2. Zero-mean noise

If, instead, we assume that the noisy-path-integration noise has zero mean then there is no systematic effect on pointing, which we demonstrate as follows for an estimate of orientation. We added a normally distributed random error to the estimate of visual direction with respect to ‘North’ on each step, *η^b^*:

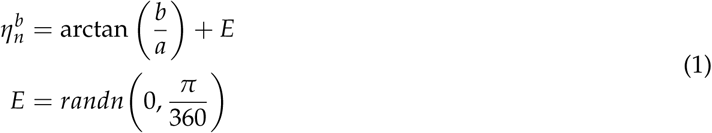

with the function *randn*(*µ*, *σ*) returning a random number drawn from a distribution with a standard deviation *σ*, and a mean *µ*. *E* is a random error added to the estimate of 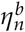, drawn from a distribution with a standard deviation of 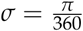 radians, and a mean of *µ* = 0.

Using Eq. (1), predictions of pointing directions were calculated with a random additive noise on the direction of ‘North’. Calculating the directions 100 times for each box in each layout, using the walking trajectory of every participant tested in the indirect walking condition of *Experiment 1* at pointing Zone C, a histogram of errors is plotted in Fig. S4d (and the same result applies for Zone A and Zone B).

#### S4.3. Abathic Model

Johnston [44] shows psychophysical data described by a linear relationship between estimated (or ‘scaling’) distance to physical viewing distance (e.g. their Fig. 7). In general, we can fit the two parameters (intercept and slope) to our pointing data (Fig. 5a). In our case, the best-fitting values are a slope of 1.03 and an intercept of 0.17, which corresponds to an abathic distance of −5.66. Specifically, the misestimated egocentric distances of the boxes, 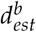 is:

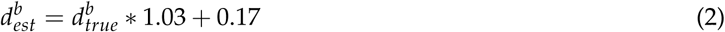

where 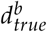 is the true egocentric distance and *b* = [1, …, 4] the index for each box.

#### S4.4. Retrofit model

We can allow box position to vary and calculate the maximum likelihood configuration of boxes that could account for the participant data (separately for each Experiment). We considered a 200 by 200 grid of possible box positions centered on the true box locations. For each possible box position the likelihood of the participant representing the box as being at that location (given their pointing responses from 3 different zones) can be defined as:

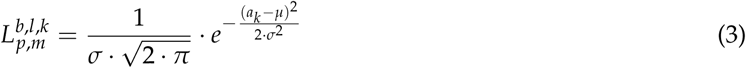

with *p* = [1, …, 20] for 20 participants, and *b* = [1, …, 4] for all 4 boxes in *l* = [1, …, 9] 9 box layouts, and *m* = [1, …, 3] for all 3 testing zones, and *k* = [1, …, 4 000] for all the possible box positions and *a_k_* is the angular error between the estimated pointing direction and the *kth* possible box position. We assume no systematic bias (*µ* = 0) and an arbitrarily chosen *σ* = 15. The maximum likelihood is then calculated for each box in each layout across all participants across all zones:

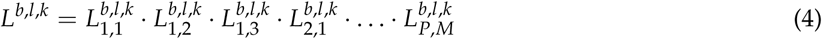

where *P* = 20 for the total number of participants and *M* = 3 for the total number of shooting zones.

**Figure S5:**
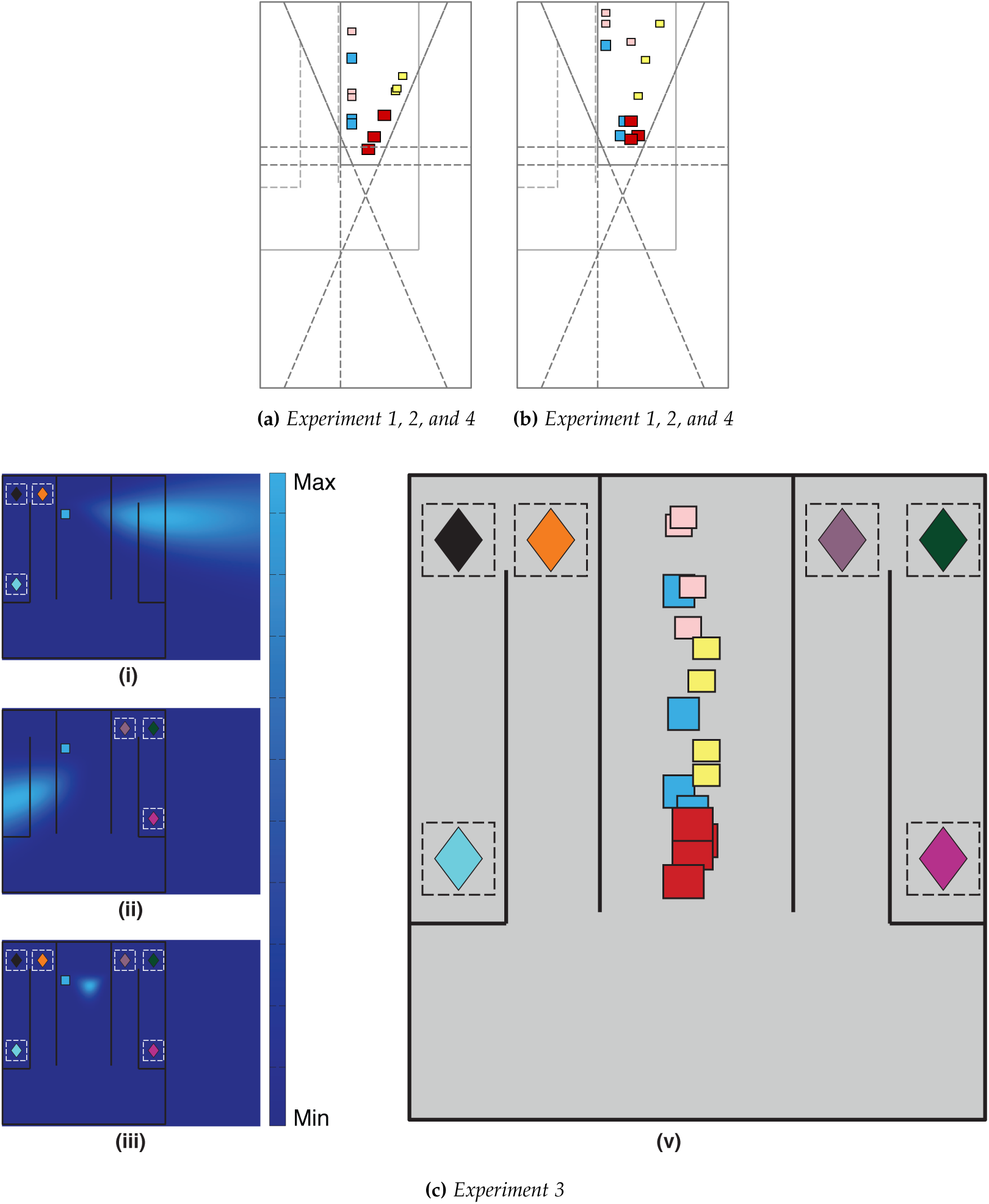
(a–b) Box layouts tested in Experiment 1, 2, and 4: (a) Real box positions of 3 box layouts plotted in the same plan view. (a) Predicted box layouts calculated using the ‘retrofit’ model, based on combined data from all studies. (c) Experiment 3, ‘retrofit’ model using the estimated directions of one participant pointing to the blue box in one layout (i) at zones A–C alone or (ii) at zones D–F alone or (iii) A–F all together. (v) Similar to (iii) but now showing the maximum likelihood location for all the boxes in all layouts. In both (iii) and (v), the predicted box locations are shifted towards a ‘North-South’ plane in the center of the room.

### S5. Model comparison

#### S5.1. Per-participant comparison between path-integration, abathic distortion and ‘projection plane’ models in Experiment 1

**Table S1:**
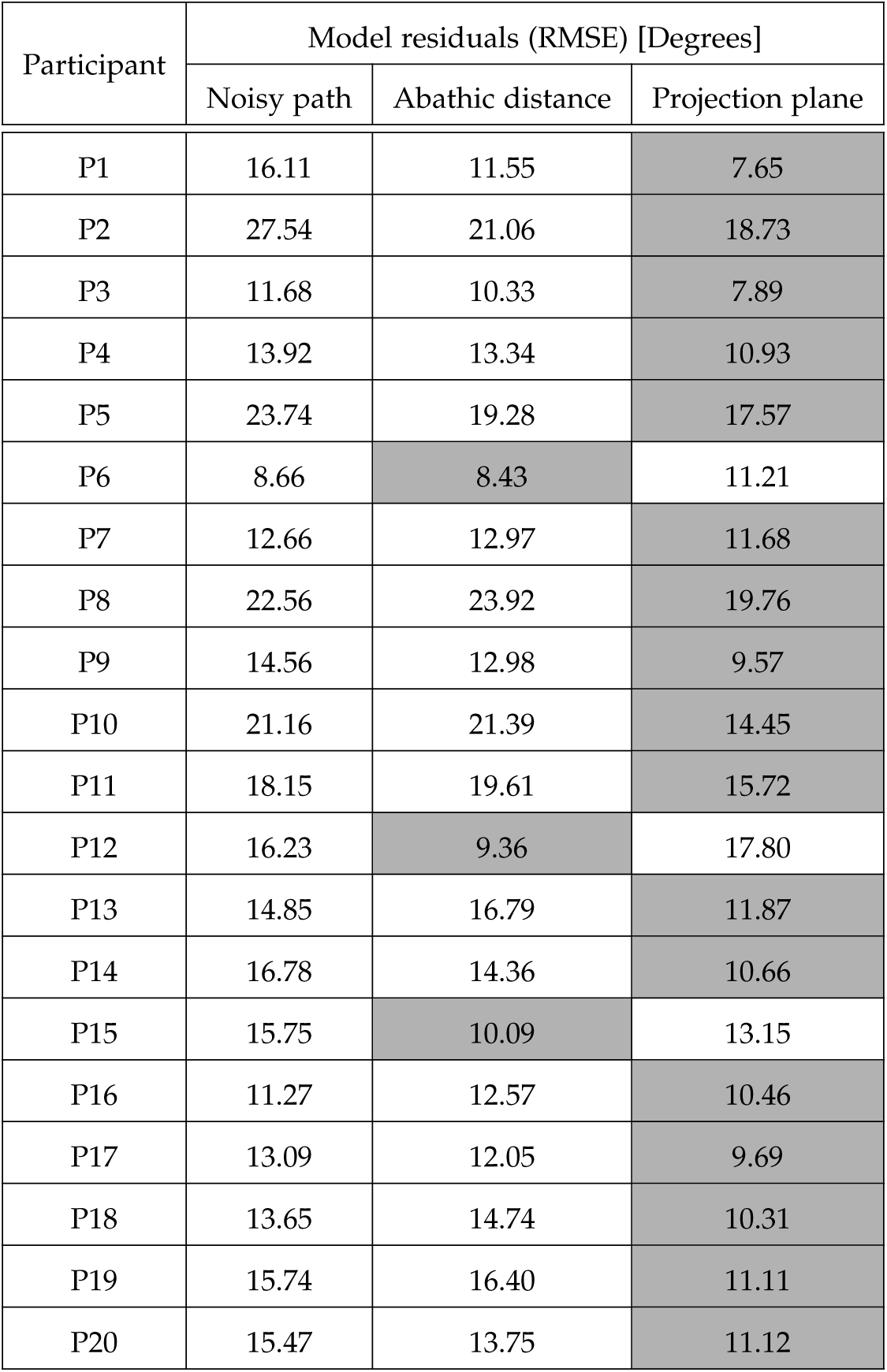
Root-mean-square residuals (RMSE) of the pointing data relative to two models of pointing error for Experiment 1. RMSE values are shown per participant for two standard models and the projection plane model. See Sections S4.1 and 3.2 for details of the noisy-path-integration and abathic models respectively. Gray-colored cells show lowest RMSE out of the 3 different models for each participant. Overall, the RMSE for the projection-plane model is 13.0, for the noisy-path-integration model is 16.8 and for the abathic-distance-distortion model is 15.3 degrees. One participant had knowledge of the type of hypothesis that was being explored in the experiments (P15). Their RMSE values in Experiment 1 shown here and for the values shown in Figure 7 and Figure 8 were within the range for other participants.

#### S5.2. Model comparisons using two model selection criteria, AIC and BIC

**Table S2:** A comparison of the fits of the projection plane model and the retrofit model using Akiake (AIC) and Bayesian (BIC) Information Criteria which penalize a model according to the number of free parameters it has. Lower values indicate a better fit.

| Experiment | 1 |  | 2 |  | 3 |  | 4 |  |
| --- | --- | --- | --- | --- | --- | --- | --- | --- |
| Criteria | Projection | Retrofit | Projection | Retrofit | Projection | Retrofit | Projection | Retrofit |
| AIC | 156.7 | 258.5 | 362.2 | 509.5 | 460.7 | 446.4 | 961.7 | 1177.7 |
| BIC | 158.3 | 298.1 | 364.5 | 566.4 | 463.3 | 510.5 | 964.9 | 1259.1 |
| # of Parameters | 0 | 24 | 0 | 24 | 0 | 24 | 0 | 24 |

### S6. Raw data

Raw data for all the figures and code to reproduce an example figure (Fig. 8) are online:

https://github.com/jnyvng/PointingData.

Also, there is an interactive website showing the raw pointing data for all the experiments:

http://www.jennyvuong.net/dataWebsite/

Videos showing what the scene in Experiment 1 looked like in the real-world experiment and in VR are available at:

https://www.glennersterlab.com/Videos/RealWorld.mp4.

https://www.glennersterlab.com/Videos/VREnv.mp4

